# Augmentation of Histone Deacetylase 6 Activity Impairs Mitochondrial Respiratory Complex I in Ischemic/Reperfused Diabetic Hearts

**DOI:** 10.1101/2023.02.21.529462

**Authors:** Shelley L. Baumgardt, Juan Fang, Xuebin Fu, Yanan Liu, Zhengyuan Xia, Ming Zhao, Ling Chen, Rachana Mishra, Muthukumar Gunasekaran, Progyaparamita Saha, Joseph M. Forbess, Zeljko J. Bosnjak, Amadou KS Camara, Judy R. Kersten, Edward Thorp, Sunjay Kaushal, Zhi-Dong Ge

**Affiliations:** Departments of Anesthesiology, Medical College of Wisconsin, 8701 Watertown Plank Road, Milwaukee, Wisconsin 53206; Cardiovascular-Thoracic Surgery and the Heart Center, Stanley Manne Children’s Research Institute, Ann & Robert H. Lurie Children’s Hospital of Chicago, Departments of Pediatrics and Surgery, Feinberg School of Medicine, Northwestern University, 225 E. Chicago Avenue, Chicago, Illinois 60611; Departments of Pathology and Pediatrics, Feinberg School of Medicine, Northwestern University, 300 E. Superior Avenue, Chicago, Illinois 60611; Department of Pediatrics, Medical College of Wisconsin, 8701 Watertown Plank Road, Milwaukee, Wisconsin 53206; Department of Anesthesiology, Affiliated Hospital of Guangdong Medical University, Zhanjiang, Guangdong Province, The People’s Republic of China; The Feinberg Cardiovascular and Renal Research Institute, Feinberg School of Medicine, Northwestern University, 300 E. Superior Avenue, Chicago, Illinois 60611; Departments of Medicine and Physiology, Medical College of Wisconsin, 8701 Watertown Plank Road, Milwaukee, Wisconsin 53206

**Keywords:** Histone deacetylase 6, Ischemia/reperfusion, Type 1 diabetes, Type 2 diabetes, Mitochondria, Tumor necrosis factor α

## Abstract

**BACKGROUND:** Diabetes augments activity of histone deacetylase 6 (HDAC6) and generation of tumor necrosis factor α (TNFα) and impairs the physiological function of mitochondrial complex I (mCI) which oxidizes reduced nicotinamide adenine dinucleotide (NADH) to nicotinamide adenine dinucleotide to sustain the tricarboxylic acid cycle and β-oxidation. Here we examined how HDAC6 regulates TNFα production, mCI activity, mitochondrial morphology and NADH levels, and cardiac function in ischemic/reperfused diabetic hearts.

**METHODS:** HDAC6 knockout, streptozotocin-induced type 1 diabetic, and obese type 2 diabetic db/db mice underwent myocardial ischemia/reperfusion injury *in vivo* or *ex vivo* in a Langendorff-perfused system. H9c2 cardiomyocytes with and without HDAC6 knockdown were subjected to hypoxia/reoxygenation injury in the presence of high glucose. We compared the activities of HDAC6 and mCI, TNFα and mitochondrial NADH levels, mitochondrial morphology, myocardial infarct size, and cardiac function between groups.

**RESULTS:** Myocardial ischemia/reperfusion injury and diabetes synergistically augmented myocardial HDCA6 activity, myocardial TNFα levels, and mitochondrial fission and inhibited mCI activity. Interestingly, neutralization of TNFα with an anti-TNFα monoclonal antibody augmented myocardial mCI activity. Importantly, genetic disruption or inhibition of HDAC6 with tubastatin A decreased TNFα levels, mitochondrial fission, and myocardial mitochondrial NADH levels in ischemic/reperfused diabetic mice, concomitant with augmented mCI activity, decreased infarct size, and ameliorated cardiac dysfunction. In H9c2 cardiomyocytes cultured in high glucose, hypoxia/reoxygenation augmented HDAC6 activity and TNFα levels and decreased mCI activity. These negative effects were blocked by HDAC6 knockdown.

**CONCLUSIONS:** Augmenting HDAC6 activity inhibits mCI activity by increasing TNFα levels in ischemic/reperfused diabetic hearts. The HDAC6 inhibitor, tubastatin A, has high therapeutic potential for acute myocardial infarction in diabetes.

**Novelty and Significance:** *What Is Known?:*

1. Ischemic heart disease (IHS) is a leading cause of death globally, and its presence in diabetic patients is a grievous combination, leading to high mortality and heart failure.
2. Diabetes impairs assembly of mitochondrial complex I (mCI), complex III dimer, and complex IV monomer into the respiratory chain supercomplexes, resulting in electron leak and the formation of reactive oxygen species (ROS).
3. By oxidizing reduced nicotinamide adenine dinucleotide (NADH) and reducing ubiquinone, mCI physiologically regenerates NAD^+^ to sustain the tricarboxylic acid cycle and β-oxidation.

*What New Information Does This Article Contribute?:*

1. Myocardial ischemia/reperfusion injury (MIRI) and diabetes as comorbidities augment myocardial HDCA6 activity and generation of tumor necrosis factor α (TNFα), which inhibit myocardial mCI activity.
2. Genetic disruption of histone deacetylase 6 (HDAC6) decreases mitochondrial NADH levels and augments mCI activity in type 1 diabetic mice undergoing MIRI via decreasing TNFα production, leading to decreases in MIRI.
3. Pretreatment of type 2 diabetic db/db mice with a HDAC6 inhibitor, tubastatin A (TSA), decreases mitochondrial NADH levels and augments mCI activity by decreasing TNFα levels, leading to improvements in cardiac function. Patients with diabetes are more susceptible to MIRI than non-diabetics with greater mortality and resultant heart failure. There is an unmet medical need in diabetic patients for the treatment of IHS. Our biochemical studies find that MIRI and diabetes synergistically augment myocardial HDAC6 activity and generation of TNFα, along with cardiac mitochondrial fission and low bioactivity of mCI. Intriguingly, genetic disruption of HDAC6 decreases the MIRI-induced increases in TNFα levels, concomitant with augmented mCI activity, decreased myocardial infarct size, and ameliorated cardiac dysfunction in T1D mice. Importantly, treatment of obese T2D db/db mice with TSA reduces the generation of TNFα and mitochondrial fission and enhances mCI activity during reperfusion after ischemia. Our isolated heart studies revealed that genetic disruption or pharmacological inhibition of HDAC6 reduces mitochondrial NADH release during ischemia and ameliorates dysfunction of diabetic hearts undergoing MIRI. Furthermore, HDAC6 knockdown in cardiomyocytes blocks high glucose- and exogenous TNFα-induced suppression of mCI activity *in vitro*, implying that HDAC6 knockdown can preserve mCI activity in high glucose and hypoxia/reoxygenation. These results demonstrate that HDAC6 is an important mediator in MIRI and cardiac function in diabetes. Selective inhibition of HDAC6 has high therapeutic potential for acute IHS in diabetes.

## INTRODUCTION

Diabetes mellitus is a growing public health problem with a prevalence approaching 422 million worldwide. Type 1 diabetes (T1D) is currently an incurable autoimmune disease marked by progressive and eventually exhaustive destruction of insulin-producing pancreatic β cells. Type 2 diabetes (T2D) is the combination of insulin resistance in peripheral tissue, insufficient insulin secretion from pancreatic β cells, and excessive glucagon secretion from pancreatic α cells. Both T1D and T2D enhance risk for ischemic heart disease (IHS) by 2- to 6-fold.^1^ Despite current optimal therapy, the mortality rate of acute myocardial infarction in diabetic patients is more than double that of nondiabetics.^2^ There is an urgent need to develop efficacious therapeutic interventions to prevent and reduce myocardial ischemia/reperfusion injury (MIRI) in diabetics.

Histone deacetylase (HDAC) 6 functions to remove acetyl groups of lysine residues from histone and nonhistone proteins.^3–5^ HDAC6-reversible lysine acetylation was recently identified as a posttranslational modification that controls mitochondrial, myofibril, sarcomere, and microtubule functions and myocardial passive stiffness.^4, 6–12^ The catalytic activity and expression of HDAC6 can be induced in response to hyperglycemic stress, MIRI, hypertension, and angiotensin II.^13–15^ Importantly, pharmacological inhibition of HDAC6 confers cardioprotection in T1D rodents with acute myocardial infarction and diabetic cardiomyopathy.^3, 16, 17^ However, little is known regarding the regulation and role of HDAC6 in ischemic/reperfused T2D hearts.

Mitochondrial complex I (mCI, NADH:ubiquinone oxidoreductase) is the first and largest enzyme complex of the mitochondrial electron transport chain.^18^ It oxidizes reduced nicotinamide adenine dinucleotide (NADH) which is generated through the Krebs cycle in the mitochondrial matrix and uses the two electrons to reduce ubiquinone to ubiquinol.^19^ Although individual mCI behaves as a single unit, mCI, complex III dimer, and complex IV monomer frequently assemble into a supramolecular unit called supercomplexes to increase the efficacy of the electron transport, reducing the rate of production of ROS.^20^ Diabetes and abnormally high levels of tumor necrosis factor α (TNFα) impair the supercomplex assembly and activity, resulting in electron leak and increased production of reactive oxygen species (ROS).^21, 22^ Emerging studies reveal during MIRI, mCI can also catalyze the reverse reaction, Δp-linked oxidation of ubiquinol to reduce NAD^+^ (or O_2_), known as reverse electron transfer (RET), to generate ROS.^23, 24^ Since mCI is susceptible to ischemic stress, it plays a seminal role in promoting oxidative injury.^23^ A proteome-wide screening study has identified lysine residues on subunits of mCI as targets of HDACs.^25^ It remains unknown whether HDAC6 affects mCI activity in ischemic/reperfused diabetic hearts.

Here, we quantitated myocardial HDAC6 activity, TNFα concentrations, and mCI activity to understand their relationship in T1D and T2D mice with and without MIRI. We then examined the effects of HDAC6 knockout (KO) on TNFα concentrations, mCI activity, mitochondrial morphology and NADH levels, and myocardial infarct size in T1D mice *in vivo* and *ex vivo*. Using a selective inhibitor of HDAC6, tubastatin A (TSA), we further investigated the effect of HDAC6 inhibition on TNFα concentrations, mCI activity, mitochondrial morphology and function, and cardiac function in T2D mice *in vivo* and *ex vivo*. These studies identify HDAC6 as a potent regulator of TNFα production and mCI activity and yield a strategy to inhibit HDAC6 to protect the heart against MIRI in diabetes. Taken together, our findings provide new insights into the pathogenesis of MIRI in diabetes and highlight opportunities to improve treatment.

## METHODS

### Data Availability

Detailed Materials and Methods are provided in the <u>Supplemental Material</u>. The data that support the findings of this study are available from the corresponding author upon reasonable request.

## RESULTS

### Cardiac HDAC6 Activity is Augmented synergistically by Diabetes and MIRI

To study the functional implications of HDAC6 in T1D and MIRI, we first examined the myocardial HDAC6 activity in T1D C57BL/6 mice with and without MIRI. Streptozotocin (STZ) is a naturally occurring alkylating antineoplastic agent that is particularly toxic to the insulin-producing β cells of the pancreas in mammals.^26^ It was used to induce T1D in C57BL/6 mice, and citrate buffer as a control (Figure S1). C57BL/6 mice with normal blood glucose levels and T1D were subjected to coronary artery occlusion for 20 min followed by reperfusion for 24 h *in vivo* or sham surgery as a control (Figure 1A). Blood glucose levels at the time point of 24 hours after post-ischemic reperfusion were comparable between T1D + MRI and T1D groups (Figure 1B). 2,3,5-Triphenyltetrazolium chloride (TTC) and phthalocyanine blue dye staining displayed that MIRI elicited a marked infarct size (infarct size/area at risk: 59 ± 3%) (Figure 1C) in T1D mice. In a separate group, mouse myocardium was collected for measuring HDAC6 activity. Compared with the Ctrl group, HDAC6 activity was significantly augmented by T1D (Figure 1D). Furthermore, it is higher in T1D + MIRI than T1D group (Figure 1D). These data indicate that T1D and MRI synergistically augment myocardial HDAC6 activity.

**Figure 1.**
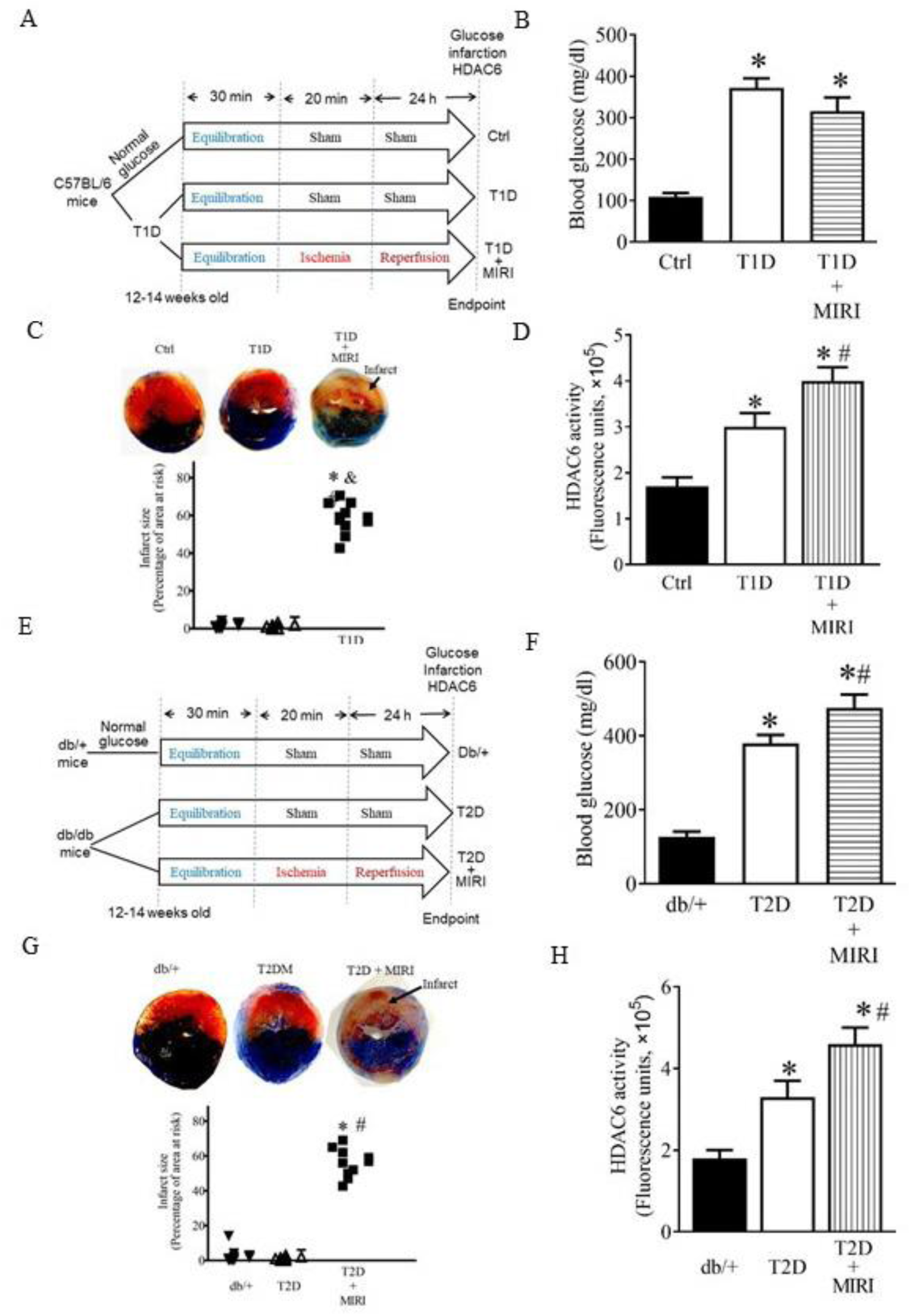
HDAC6 activity was augmented synergistically by diabetes and MIRI. A: experimental procedures for STZ-induced T1D mice; B: fasting blood glucose in T1D and Ctrl mice 24 h after ischemia or sham surgery; C: top: representative heart sections stained with TTC and phthalocyanine blue showing area at risk (red + white) and infarct size (white). Bottom: infarct size expressed as a percentage of area at risk in T1D and Ctrl mice; D: Myocardial HDAC6 activity of T1D, T1D + MIRI, and Ctrl mice. *P < 0.05 versus Ctrl and ^#^P < 0.05 versus T1D (n = 9-10 mice/group). E: experimental procedures for obese type 2 diabetic (T2D) db/db mice and controls (db/+); F: fasting blood glucose in db/db mice and Ctrl 24 h after ischemia or sham surgery; G: top: heart sections stained with TTC and phthalocyanine blue showing area at risk and infarct size. Bottom: infarct size expressed as a percentage of area at risk in T2D and Ctrl mice; H: myocardial HDAC6 activity of T2D, T2D + MIRI, and db/+ mice. *P < 0.05 versus db/+ and ^#^P < 0.05 versus T2D (n = 9-10 mice/group).

We next determined myocardial HDAC6 activity in obese T2D mice with and without MIRI. Leptin receptor-deficient homozygous db mice, *db*/*db*, are among the most widely used animal models in obesity-induced T2D.^27^ Obese T2D db/db and non-obese heterozygous db/+ Ctrl mice (Figure S2) underwent coronary artery occlusion for 20 min followed by reperfusion 24 hours or sham surgery as a control (Figure 1E). Blood glucose levels were higher in T2D than db/+ groups and in T2D + MIRI than T2D groups (Figure 1F). Coronary artery occlusion for 20 min followed by reperfusion for 24 hours elicited 53 ± 4% of infarct size/area-at-risk (Figure 1G). Myocardial HDAC6 activity was higher in T2D than db/+ groups and in T2D + MIRI than T2D groups (Figure 1H). These data suggest that T2D and MIRI synergistically augment myocardial HDAC6 activity.

### Mitochondrial Morphology and mCI are Impaired Jointly by Diabetes and MIRI

To investigate mitochondrial damage in T1D and MIRI, we used transmission electron microscopy (TEM) to visualize subcellular structures and organization of the myocardium with a focus on the mitochondrial. In the Ctrl mice, mitochondria and myofilaments were orderly arranged, and no fusion was observed between mitochondrial cristae and membranes (Figure 2A). In the T1D group, myofilaments and mitochondria were disordered, mitochondria were swollen and showed great fission, and mitochondrial cristae and membranes were dissolved and changes in the vacuole. Compared to the T1D group, the mitochondria in the T1D + MIRI group were highly disordered, exhibiting more edema and dissolution with mitochondrial cristae and membrane rupture. We quantified mitochondrial volume density and mitochondrial surface area in 20-21 sections from 3 mice/group. There were no significant differences in mitochondrial volume density among 3 groups (p > 0.05) (Figure 2A). Mitochondrial surface area was smaller (mitochondrial fission) in T1D than Ctrl groups and in T1D + MIRI than T1D groups. These data demonstrate that T1D and MIRI jointly increase mitochondrial fission.

**Figure 2.**
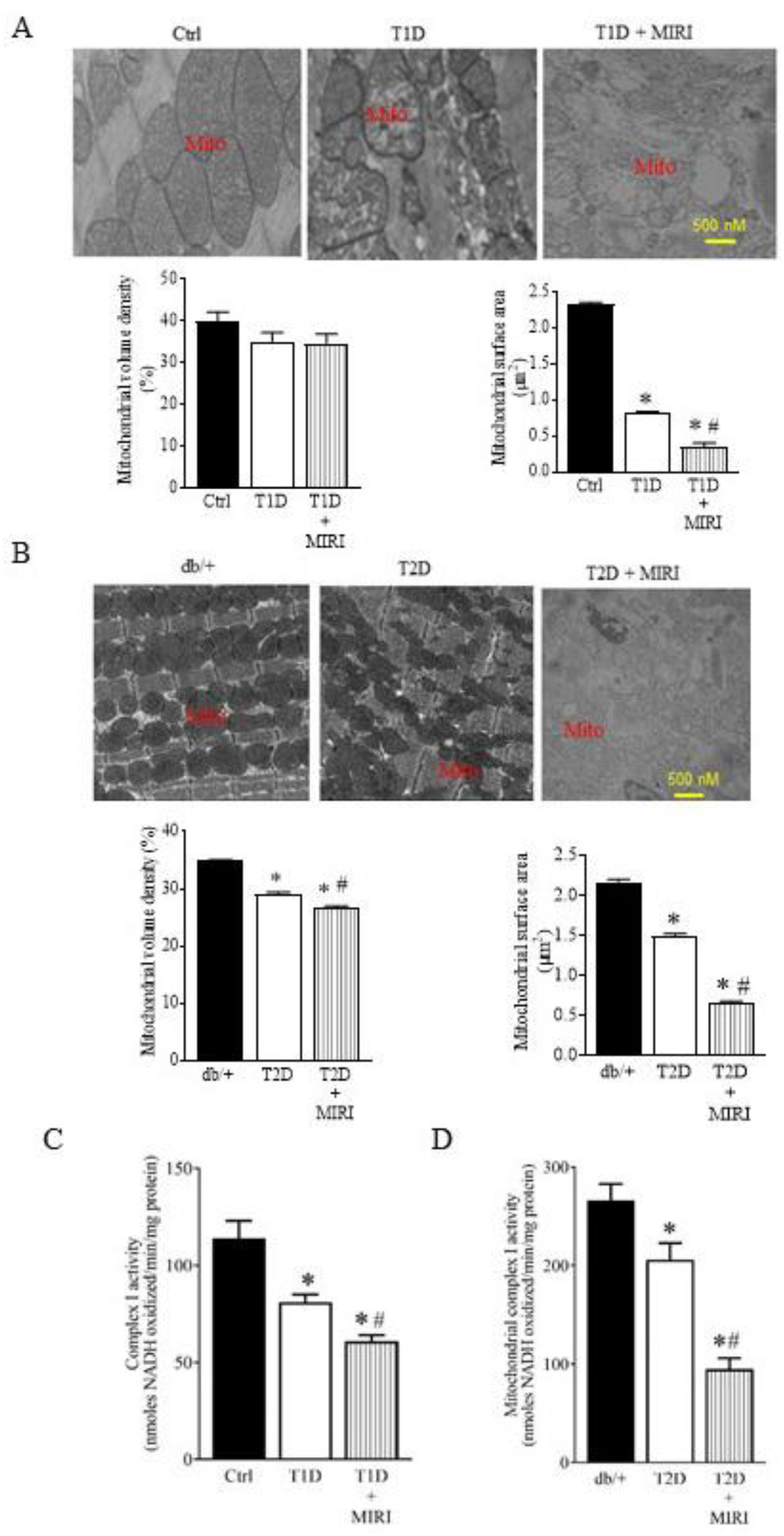
Mitochondrial morphology and mCI activity were impaired by T1D, T2D, and MIRI. A: Top: representative electron microscope micrographs showing the changes in mitochondria of T1D mice and controls. Bottom: mitochondrial volume density (n = 20-21 sections from 3 mice/group) and mitochondrial surface area (n = 200-210 mitochoondria from 3 mice/group). Ctrl, non-diabetic C57BL/6 mice were subjected to sham surgery; T1D, type 1 diabetic C57BL/6 mice underwent sham surgery; T1D + MIRI; diabetic C57BL/6 mice underwent 20 min of ischemia followed by reperfusion for 24 hours. B: Top: representative electron microscope micrographs showing the changes in mitochondria of T2D mice and controls. Bottom: mitochondrial volume density (n = 20-21 sections from 3 mice/group) and mitochondrial surface area (n = 200-210 mitochoondria from 3 mice /group). Db/+, non-diabetic db/+ mice were subjected to sham surgery; T2D, type 2 diabetic db/db mice underwent sham surgery; T2D + MIRI; db/db mice underwent 20 min of ischemia followed by reperfusion for 24 hours. C: myocardial mCI activity 5 min after post-ischemic reperfusion (n=6-8 hearts/group) in T1D mice and Ctrl mice. *P<0.05 versus Ctrl, ^#^P < 0.05 versus T1D groups. D: myocardial mCI activity 5 min after reperfusion (n=6-8 hearts/group) in T2D mice and Ctrl mice. Scale bar: 500 nm.

We next examined how T2D or MIRI alone or in combination affected mitochondrial morphology. The mitochondria of the T2D mice undergoing coronary artery occlusion for 20 min followed by reperfusion for 24 hours or sham surgery were imaged with a TEM. Heterozygous db/+ mice that had undergone sham surgery were used as a Ctrl (Figure 1E). The mitochondria in the Ctrl group were well organized, whereas mitochondria in the db/db mice that had undergone sham surgery (T2D) were fragmented (Figure 2B). Mitochondrial volume density was significantly lower in T2D than Ctrl groups and in T2D + MIRI than T2D groups, suggesting that that cardiac mitochondrial biogenesis is suppressed in the db/db mice (P < 0.05, n = 20-21 from 3 mice/group) (Figure 2B). Moreover, mitochondrial surface area was significantly smaller in T2D than Ctrl groups and in T2D + MIRI than T2D groups. These results suggest that T2D and MIRI jointly cause mitochondrial fission.

To measure mCI activity in diabetes with and without MIRI, we isolated mitochondria from mouse hearts (Figure S3) at the time point of 5 min after reperfusion in T1D and T2D mice. mCI activity was significantly lower in T1D than Ctrl groups and in T1D + MIRI than T1D groups (P < 0.05, n = 6-8 mice/group) (Figure 2C). Like the results of T1D, mCI activity was significantly lower in T2D group than db/+ group and further lower in T2D + MIRI group compared with T2D group (P < 0.05 between T2D and T2D + MIRI groups). These results indicate that diabetes and MIRI jointly inhibit myocardial mCI activity.

### Abnormal High Levels of TNFα Inhibit mCI Activity in Ischemic/Reperfused Diabetic Hearts

TNFα is a multifunctional proinflammatory cytokine produced by monocytes, macrophages, neutrophils, CD4^+^ T cells, and cardiomyocytes.^28–30^ The inhibitory effect of TNFα on the mitochondrial electron transport chain occurs in the early stage of mitochondrial damage without any pronounced damage to other cellular organells.^21^ To investigate the association of TNFα levels with low mCI activity in ischemic/reperfused T1D hearts, we used a rat IgG1 anti-TNFα monoclonal antibody, MP6-XT22, to neutralize TNFα *in vivo* (Figure 3A). Rat IgG1 monoclonal antibody, GL1B, was used as a control. MP6-XT22 significantly decreased myocardial and plasma TNFα levels but did not change mCI activity in sham-operated non-diabetic mice (P < 0.05 between Anti-TNFα mAB and Ctrl groups) (Figures 3B, 3C, and 3D). Myocardial and plasma TNFα levels were significantly increased, and mCI activity was decreased by T1D alone and in combination with MIRI. Intriguingly, MP6-XT22 significantly decreaseed myocardial and plasma TNFα levels and augmented myocardial mCI activity (P < 0.05 between T1D + Anti-TNFα mAB and T1D groups and between T1D + MIRI + Anti-TNFα mAB and T1D + MIRI groups) in ischemic/reperfused diabetic hearts (Figures 3B, 3C, and 3D). These data suggest that increased levels of TNFα contribute to low mCI activity in T1D mice undergoing MIRI.

**Figure 3.**
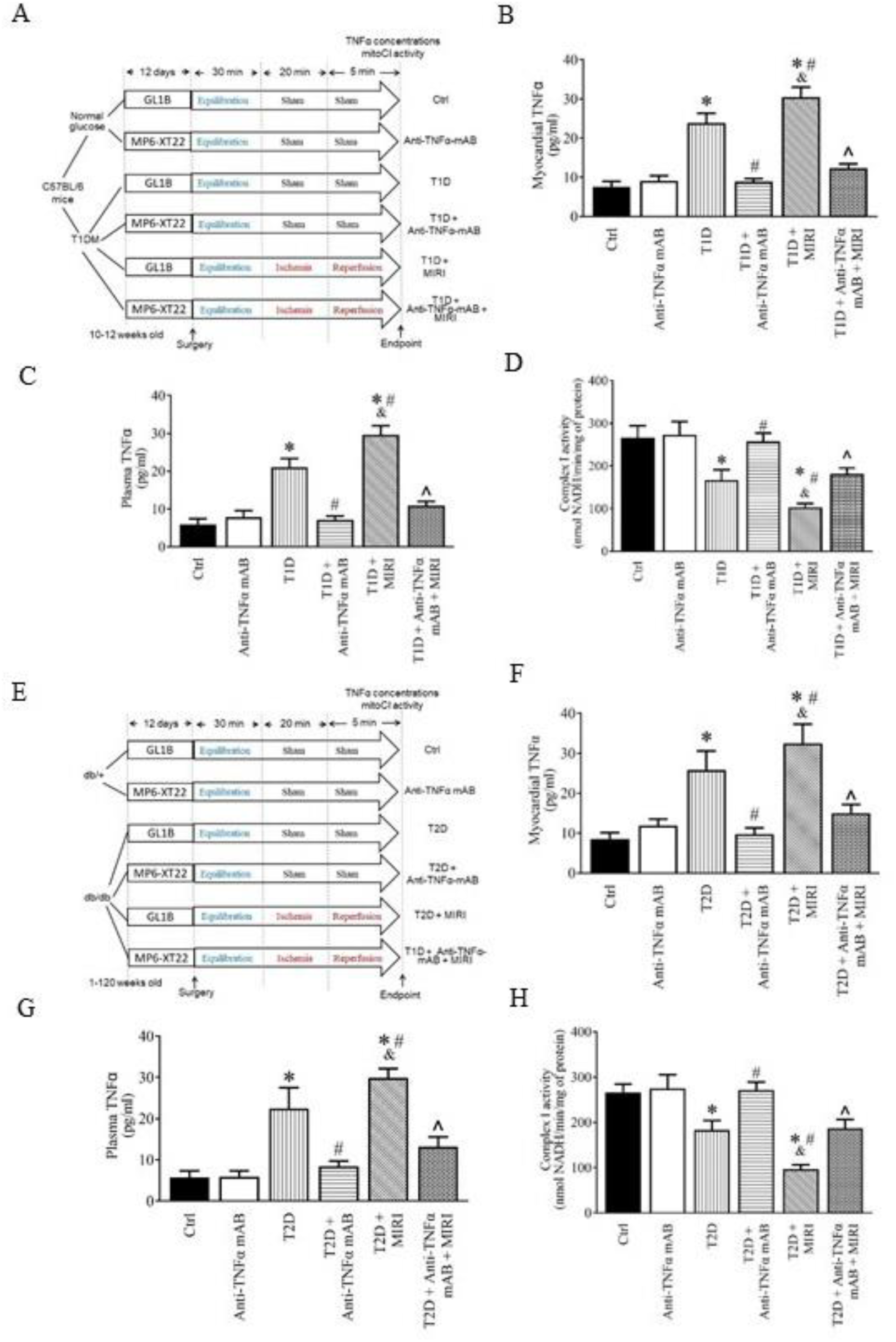
Neutralization of TNFα blocked both T1D and T2D-induced suppression of mCI activity following MIRI. A: schematic representation of the experimental procedures for neutralization of TNFα with an anti-TNFα monoclonal antibody, MP6-XT22, in mice of T1D undergoing MIRI. GL1B was used as control for MP6-XT22; B: myocardial TNFα levels in T1D and non-diabetic C57BL/6 mice 5 min after reperfusion; C: plasma TNFα levels in T1D and non-diabetic C57BL/6 mice 5 min after reperfusion; D: myocardial mCI activity 5 min after reperfusion (n=6-8 hearts/group). *P<0.05 versus Ctrl and anti-TNFα mAB group, ^#^P < 0.05 versus T1D group, ^&^P<0.05 versus T1D + anti-TNFα mAB group, and ^^^P<0.05 versus T1D + MIRI group. E: experimental procedures for the neutralization of TNFα with MP6-XT22 in T2D db/db mice undergoing MIRI; F: myocardial TNFα levels in T2D and non-diabetic db/+ mice 5 min after reperfusion; G: plasma TNFα levels in T2D and non-diabetic db/+ mice 5 min after reperfusion; H: myocardial mCI activity 5 min after reperfusion (n=6-8 hearts/group). *P<0.05 versus Ctrl gropup, *P<0.05 versus Ctrl and anti-TNFα mAB group, ^#^P < 0.05 versus T1D group, ^&^P<0.05 versus T1D + anti-TNFα mAB group, and ^^^P<0.05 versus T1D + MIRI group.

We next investigated the effects of TNFα neutralization on mCI activity in T2D mice with and without MIRI. Db/db and db/+ Ctrl mice were treated with MP6-XT22 or GL1B for 12 days prior to MIRI or sham surgery (Figure 3E). Increased levels of myocardial and plasma TNFα in T2D mice undergoing MIRI or sham surgery were significantly decreased by anti-TNFα mAB (P< 0.05 between T2D + anti-TNFα mAB and T2D groups and between T2D + MIRI + anti-TNFα mAB and T2D + MIRI groups) (Figures 3F and 3G). Interestingly, myocardial mCI activity was significantly augmented by anti-TNFα mAB in T2D undergoing MIRI (P < 0.05 between T2D + MIRI + anti-TNFα mAB and T2D + MIRI groups) (Figure 3H). These results suggest that increased levels of TNFα are responsible for inhibition of mCI activity in T2D mice subjected to MIRI.

### Genetic Disrupion of HDAC6 Reduces TNFα Levels and Augments mCI Acvitity in Ischemic/Reperfused T1D Mice

Since myocardial HDAC6 activity is greatly augmented in ischemic/reperfused T1D rats,^13^ we investigated whether HDAC6 knockout (KO) affected blood glucose and plasma TNFα levels, myocardial infarct size, and mCI activity. HDAC6^-/-^ and C57BL/6 mice were intraperitoneally injected STZ to induce T1D or citrate buffer as a control (Figure 4A). Supplementary Table 1 lists the general characteristics and echocardiographic parameters (Figure S4) of 4 groups of mice: MIRI, HDAC6^-/-^ + MIRI, T1D + MIRI, HDAC6^-/-^ + T1D + MIRI, prior to MIRI surgery. Fasting blood glucose levels at the time point of 24 hours after post-ischemic reperfusion were comparable between HDAC6^-/-^ + MIRI and MIRI groups but were higher in T1D + MIRI than MIRI and HDAC6^-/-^ + MIRI groups (Figure 4B). Compared to the T1D + MIRI group, blood glucose levels were decreased in HDAC6^-/-^ + T1D + MIRI group. We determined plasma TNFα levels and myocardial mCI activity 5 min after post-ischemic reperfusion in HDAC6^-/-^ and C57BL/6 mice with and without T1D (Figure 4A). Both plasma TNFα levels and mCI activity were comparable between HDAC6^-/-^ + MIRI and MIRI groups. Compared to the MIRI or HDAC6^-/-^ + MIRI groups, plasma TNFα levels were significantly increased (Figure 4C), and mCI activity was inhibited in T1D + MIRI groups (P < 0.05, n = 9-11 mice/group) (Figure 4D). Interestingly, plasma TNFα levels were lower, and mCI activity was higher in HDAC6^-/-^ + T1D + MIRI than T1D + MIRI groups. These results suggest that HDAC6 KO decreases plasma TNFα levels and augments myocardial mCI activity in ischemic/reperfused T1D mice.

**Supplementary Table 1.** General characteristics and echocardiographic parameters of C57BL/6 and HDAC6^-/-^ mice prior to myocardial ischemia/reperfusion injury.

|  | MIRI | HDAC6<sup>-/-</sup><br>+ MIRI | T1D<br>+ MIRI | HDAC6<sup>-/-</sup><br>+ T1D + MIRI |
| --- | --- | --- | --- | --- |
| Body weight, g | 28.3±0.3 | 28.1±0.2 | 22.7±0.3** | 24.2±0.3**^ |
| Heart rate, beats/min | 477±16 | 470±19 | 452±15 | 446±16 |
| Blood glucose, mg/dl | 127±9 | 113±11 | 338±15** | 349±16** |
| Myocardial HDAC6 activity,<br>fluorescence units, ×10<sup>5</sup> | 1.7±0.2 | 1.3±0.2 | 3.0±0.3** | 2.0±0.2^ |
| Plasma TNFα, pg/ml | 6.8±1.7 | 6.0±1.7 | 12.1±1.5** | 7.7±1.1^ |
| MAP, mmHg | 103±4 | 107±3 | 99±2 | 106±3 |
| SAP, mmHg | 137±5 | 140±4 | 129±4 | 138±4 |
| DAP, mmHg | 87±5 | 90±3 | 84±2 | 89±3 |
| Anterior wall at end diastole, mm | 1.01±0.08 | 1.05±0.08 | 0.87±0.03 | 0.99±0.05 |
| Anterior wall at end systole, mm | 1.51±0.10 | 1.47±0.11 | 1.31±0.06 | 1.43±0.09 |
| Posterior wall at end diastole, mm | 1.10±0.07 | 1.02±0.06 | 0.95±0.03 | 0.95±0.04 |
| Posterior wall at end systole, mm | 1.56±0.08 | 1.62±0.11 | 1.46±0.09 | 1.48±0.07 |
| Left ventricular end-diastolic volume, μl | 64±5 | 67±5 | 71±5 | 67±5 |
| Left ventricular end-systolic volume, μl | 17±2 | 17±2 | 20±1 | 20±3 |
| Ejection fraction, % | 74±3 | 75±2 | 70±2 | 71±3 |
| Cardiac output, ml/min | 19±2 | 21±2 | 20±1 | 21±2 |
| Mitral E/e' ratio | 22±1 | 23±1 | 24±3 | 23±2 |

DAP, diastolic pressure; MAP, mean arterial pressure; MIRI, myocardial ischemia/reperfusion injury; SAP, systolic pressure; T1D, type 1 diabetes. \*P<0.05 versus MIRI groups, #P<0.05 versus HDAC6<sup>-/-</sup> + MIRI groups, ^P<0.05 versus T1D + MIRI groups (n = 10-12 mice/group).

**Figure 4.**
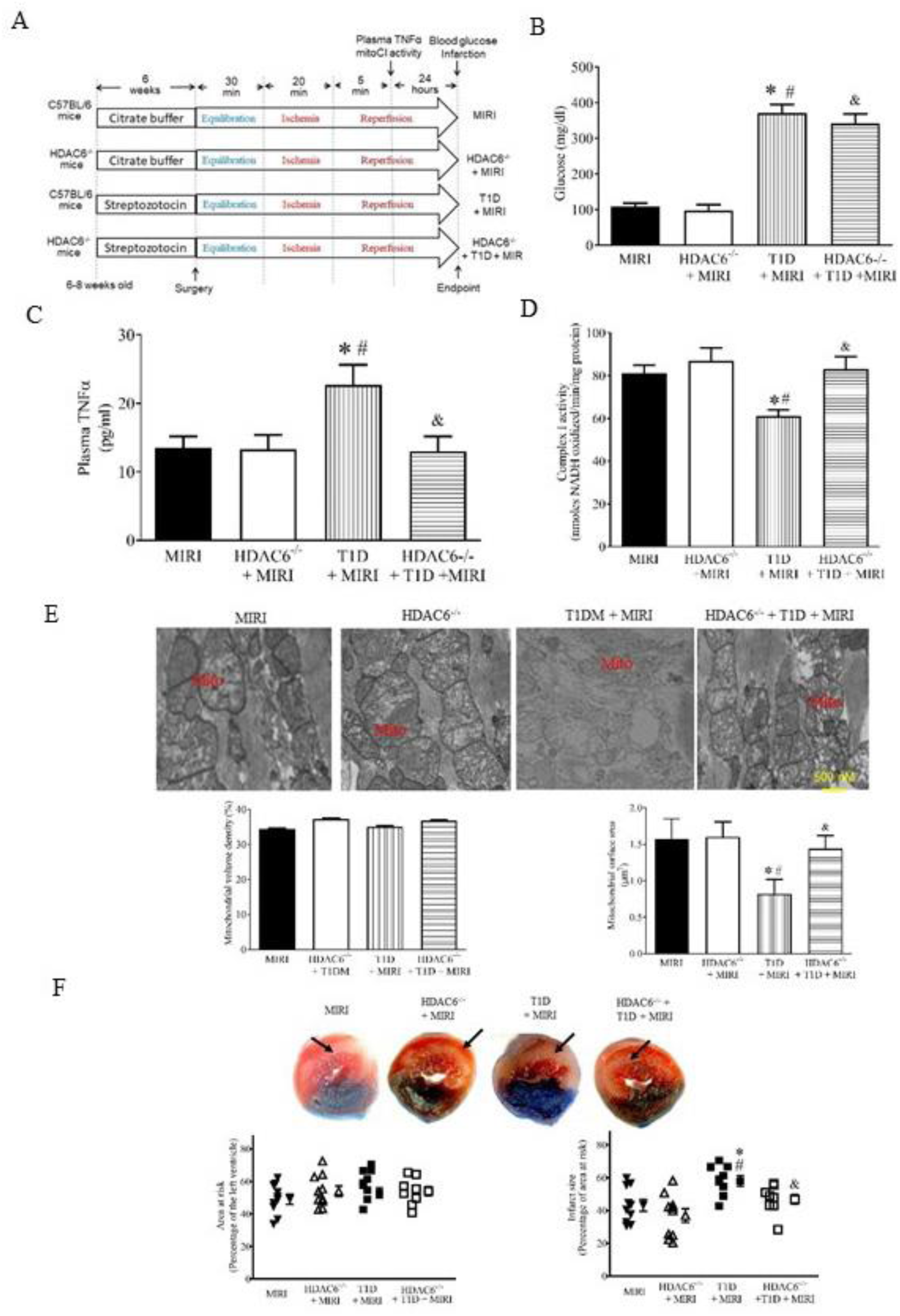
HDAC6 knockout decreased plasma TNFα and infarct size and augmented mCI activity during reperfusion in type 1 diabetic mice. A: experimental procedures; B: fasting blood glucose in HDAC6^-/-^ and C57BL/6 mice; C: plasma TNFα levels in HDAC6^-/-^ and C57BL/6 mice; D: mCI activity (n = 9-11 hearts/group); E: top: representative electron microscope micrographs of mitochondria. Bottom: mitochondrial volume density (n = 20-21 sections from 3 mice/group) and mitochondria surface area (n = 200-210 mitochondria from 3 mice/group); F: top: representative heart images showing area at risk (non-black area) and infarct size. Arrows point to infarct area (white). Botom: area at risk expressed as a percentage of the left ventricle and infarct size expressed as a percentage of area at risk (n = 8-10 mice/group). Data are presented as means ± SEM. Kruskal-Wallis test followed by Dunn’s test was used to analyze multiple group comparisons. *P < 0.05 versus MIRI groups, ^#^P < 0.05 versus HDAC6^-/-^ + MIRI groups, ^&^P < 0.05 versus T1D + MIRI groups.

Mitochondrial morphology affects suceptibility of the heart to MIRI, and inhibiting mitochondrial fission protects the heart against MIRI.^31^ We next used TEM to visualize cardiac mitochondria and quantify mitochondrial volume density and mitral surface area in HDAC6^-/-^ and C57BL/6 mice with and without T1D. There were no significant differences in mitochondrial volume density among the 4 groups (P > 0.05, n = 20-21 sections from 3 hearts/group) (Figure 4E). Surface area of cardiac mitochondria was comparable between HDAC6^-/-^ + MIRI and MIRI groups but was significantly smaller in T1D + MIRI than MIRI and HDAC6^-/-^ + MIRI groups (P < 0.05, n = 200-210 mitochondria from 3 hearts/group). Interestingly, mitochondrial surface area was greater in HDAC6^-/-^ + T1D +MIRI than T1D + MIRI groups. These results demonstrate that HDAC6 KO prevents mitochondrial fission during MIRI in T1D.

In separate experiments, we used phthalocyanine blue and TTC to stain mouse hearts to delineate area at risk and infarct size, respectively.^32^ Area at risk was comparatable among the 4 experimental groups (P > 0.05). Compared to the MIRI groups, infarct size was not significantly altered in HDAC6^-/-^ + MIRI group but increased in T1D + MIRI group (Fgiure 4F) (P < 0.05, n = 8-10 mice/group). Interstingly, infarct size was significantly decreased in HDAC6^-/-^ + T1D + MIRI group compared to the T1D + MIRI group. These results indicate that HDAC6 KO protects T1D hearts from MIRI.

### Pharmacological Inhibition of HDAC6 Reduces TNFα Levels and Augments mCI Activity in Ischemic/Reperfused T2D Mice

We further explored whether HDAC6 inhibition affected plasma TNF levels and myocardial mCI activity in T2D mice that had undergone MIRI. Tubastatin A (TSA) is a potent and highly specific HDAC6 inhibitor with an IC_50_ of 15 nM and more than 1000-fold selectivity toward all other isoforms except HDAC8 (57-fold).^33^ db/db and db/+ mice were pretreated with TSA 30 min before surgery for MIRI or vehicle as a control (Figure 5A). Supplementary Table 2 lists the general characteristics and echocardiographic parameters of 4 groups of mice: MIRI, TSA + MIRI, T2D + MIRI, TSA + T2D + MIRI, prior to MIRI surgery. Fasting blood glucose levels were comparable between TSA + MIRI and MIRI groups and higher in both T2D + MIRI and TSA+ T2D + MIRI groups than both MIRI and TSA + MIRI groups (Figure 5B). There were no significant differences in blood glucose levels between TSA + T2D + MIRI and T2D + MIRI groups. At the time point of 5 min post-ischemic reperfusion, plasma TNFα levels were significantly lower, and myocardial mCI activity was higher in TSA + MIRI than MIRI groups (Figures 5C and 5D). Compared to the MIRI and TSA + MIRI groups, plasma TNFα levels were enhanced, and myocardial mCI activity was inhibited in T2D +MIRI groups. Interestingly, TSA treatment significantly decreased plasma TNFα levels and augmented mCI activity in TSA + T2D + MIRI groups (P < 0.05, n = 9-11 mice/group). These data suggest that TSA decreases plasma TNFα and augments mCI activity in the ischemic/reperfused T2D mice.

**Supplementary Table 2.**
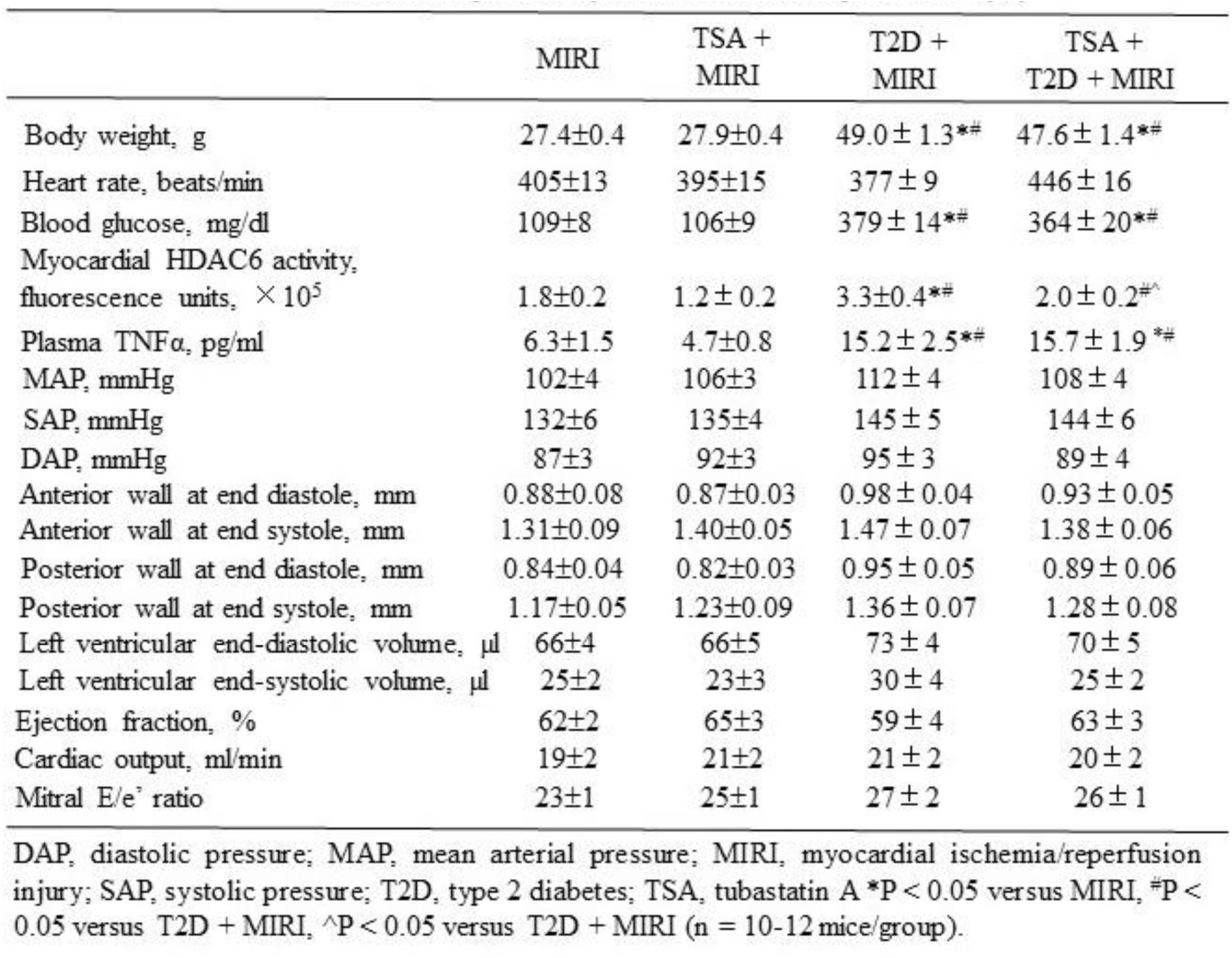
General characteristics and echocardiographic parameters of db/+ and db/db mice prior to myocardial ischemia/reperfusion injury.

|  | MIRI | TSA +<br>MIRI | T2D +<br>MIRI | TSA +<br>T2D + MIRI |
| --- | --- | --- | --- | --- |
| Body weight, g | 27.4±0.4 | 27.9±0.4 | 49.0± 1.3** | 47.6± 1.4** |
| Heart rate, beats/min | 405±13 | 395±15 | 377± 9 | 446± 16 |
| Blood glucose, mg/dl | 109±8 | 106±9 | 379± 14** | 364± 20** |
| Myocardial HDAC6 activity,<br>fluorescence units, $\times 10^5$ | 1.8±0.2 | 1.2± 0.2 | 3.3±0.4** | 2.0± 0.2<sup>^</sup> |
| Plasma TNFα, pg/ml | 6.3±1.5 | 4.7±0.8 | 15.2± 2.5** | 15.7± 1.9<sup>^</sup> |
| MAP, mmHg | 102±4 | 106±3 | 112± 4 | 108± 4 |
| SAP, mmHg | 132±6 | 135±4 | 145± 5 | 144± 6 |
| DAP, mmHg | 87±3 | 92±3 | 95± 3 | 89± 4 |
| Anterior wall at end diastole, mm | 0.88±0.08 | 0.87±0.03 | 0.98± 0.04 | 0.93± 0.05 |
| Anterior wall at end systole, mm | 1.31±0.09 | 1.40±0.05 | 1.47± 0.07 | 1.38± 0.06 |
| Posterior wall at end diastole, mm | 0.84±0.04 | 0.82±0.03 | 0.95± 0.05 | 0.89± 0.06 |
| Posterior wall at end systole, mm | 1.17±0.05 | 1.23±0.09 | 1.36± 0.07 | 1.28± 0.08 |
| Left ventricular end-diastolic volume, μl | 66±4 | 66±5 | 73± 4 | 70± 5 |
| Left ventricular end-systolic volume, μl | 25±2 | 23±3 | 30± 4 | 25± 2 |
| Ejection fraction, % | 62±2 | 65±3 | 59± 4 | 63± 3 |
| Cardiac output, ml/min | 19±2 | 21±2 | 21± 2 | 20± 2 |
| Mitral E/e' ratio | 23±1 | 25±1 | 27± 2 | 26± 1 |

DAP, diastolic pressure; MAP, mean arterial pressure; MIRI, myocardial ischemia/reperfusion injury; SAP, systolic pressure; T2D, type 2 diabetes; TSA, tubastatin A \* $P < 0.05$ versus MIRI, <sup>^</sup> $P < 0.05$ versus T2D + MIRI, <sup>^</sup> $P < 0.05$ versus T2D + MIRI ( $n = 10-12$ mice/group).

**Figure 5.**
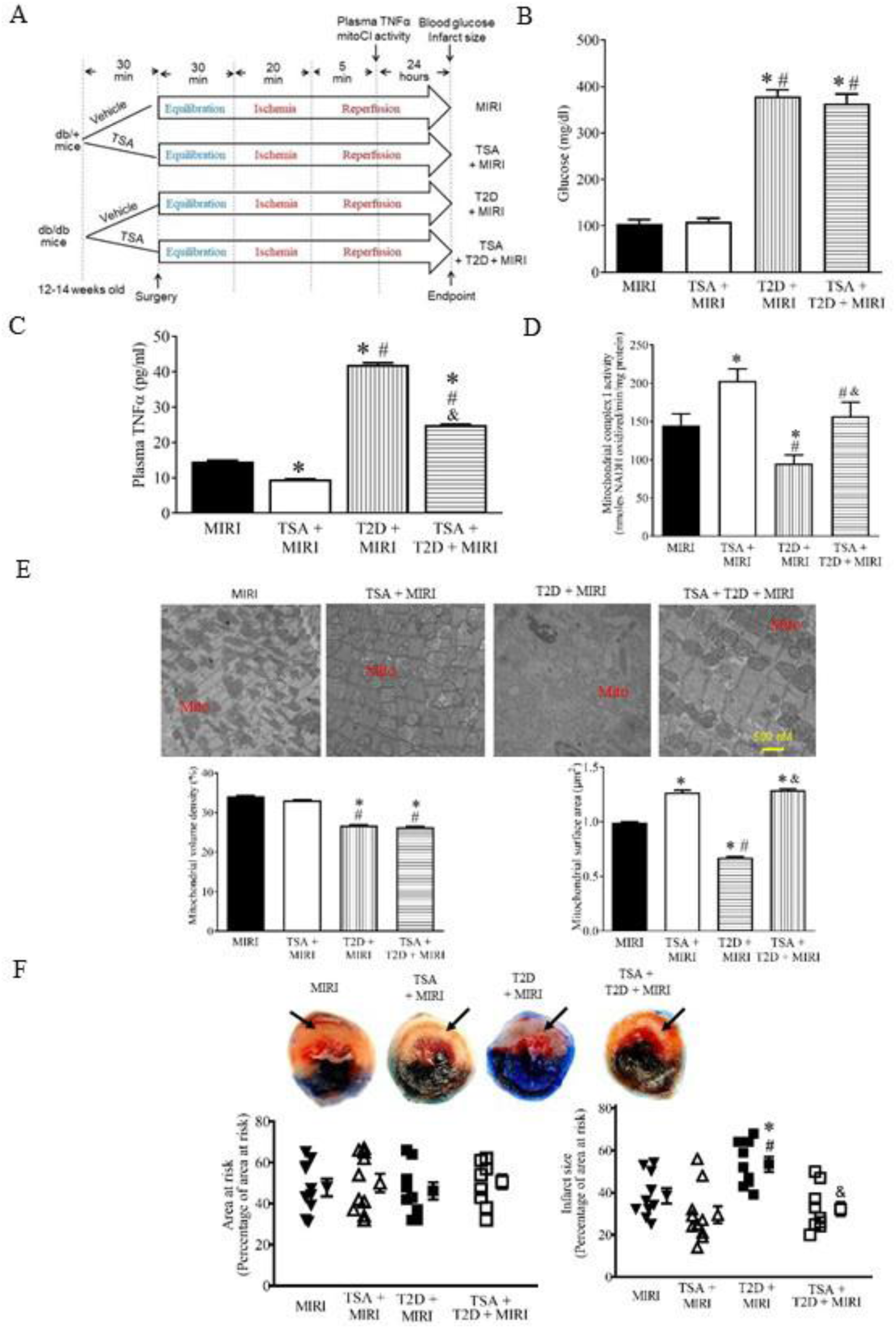
Inhibition of HDAC6 decreased plasma TNFα and infarct size and augmented mCI activity during reperfusion in T1D mice. A: experimental procedures; B: fasting blood glucose in db/db mice and db/+ control mice; C: plasma TNFα levels in db/db mice and db/+ control mice; D: myocardial mCI activity; E: top: representative electron microscope micrographs of mitochondria. Bottom: mitochondrial volume density (n = 20-21 sections from 3 mice/group) and mitochondria surface area (n = 200-210 mitochondria from 3 mice/group); F: top: representative heart images showing area at risk (non-black area) and infarct size. Arrows point to infarct area (white). Botom: area at risk expressed as a percentage of the left ventricle and infarct size expressed as a percentage of area at risk (n = 8-10 mice/group). *P < 0.05 versus MIRI groups, ^#^P < 0.05 versus TSA + MIRI groups, ^&^P < 0.05 versus T2D + MIRI groups.

The myocardium at 24 hours after post-ischemic reperfusion was imaged with a TEM to visualize mitochondria (Figure 5E). In the MIRI and TSA + MIRI groups, myofilaments and mitochondria were disordered, mitochondria were swollen and showed great fission, and mitochondrial cristae and membranes were dissolved and changes in the vacuole. Compared with the MIRI group, the mitochondria in the T2D + MIRI group were highly disordered, exhibiting more edema and dissolution with mitochondrial cristae and membrane rupture. In contrast, the hearts of the TSA + T2D + MIRI group had preserved mitochondrial morphology, exhibiting moderate edema and cristae dissolution. We quantified mitochondrial volume density and mitochondrial surface area from 20-21 sections of 3 mice in each group. Mitochondrial volume density was comparable between TSA + MIRI and MIRI groups. Compared with MIRI and TSA + MIRI groups, it was significantly decreased in T2D + MIRI and TSA + T2D + MIRI groups. There were no significant differences in mitochondrial volume density between TSA + MIRI groups and between TSA + T2D + MRI and T2D + MIRI groups (Figure 5E). Mitochondrial surface area was greater in TSA + MIRI than MIRI groups and smaller in T2D + MIRI than MIRI and TSA + MIRI groups. Interestingly, compared with T2D + MIRI group, mitochondrial surface area was significantly increased in TSA + T2D + MIRI group. These results indicate that HDAC6 inhibition prevents cardiac mitochondrial fission in ischemic/reperfused T2D mice.

The hearts were stained with phthalocyanine blue dye and TTC to delineate area at risk and infarct size, respectively (Figure 5F). There were no significant differences in area at risk among the 4 experimental groups. Coronary occlusion for 20 min followed by reperfusion for 24 h resulted in infarct size of 39 ± 4% of area at risk (n = 10 mice) in Ctrl mice, which significantly increased to 53 ± 4% (n = 10 mice, P < 0.05 versus Ctrl groups) in T2D group (Figure 5F). TSA treatment did not significantly change infarct size in db/+ mice. Interestingly, T2D-induced increase in infarct size was blocked by TSA (n = 9 mice, P < 0.05 versus T2D) (Figure 5F). These results indicate that selective inhibition of HDAC6 protects T2D mice from MIRI.

### Genetic Disruption of HDAC6 Limits Mitochondrial NADH Release and Ameliorats Cardiac Dysfunction in Ischemic/Reperfused T1D Mice

mCI oxidizes reduced form of nicotinamide adenine dinucleotide (NADH) from the tricarboxylic acid cycle and β oxidation of fatty acids, reduces ubiquinone and transports protons across the inner membrane, contributing to the proton-motive force.^34^ Since HDAC6 KO augments mCI activity in ischemic/reperfused T1D hearts, we dynamically determined the levels of NADH online during ischemia and reperfusion in isolated Langendorff-perfused hearts.^32^ Langendorff-perfused hearts were subjected to global ischemia for 30 min followed by reperfusion for 120 min (Figure 6A). Mitochondrial NADH levels from Langendorff-perfused hearts at baseline were higher in T1D and HDAC6^-/-^ + T1D groups than MIRI groups (P < 0.05, n = 8 hearts/group) (Figure 6B). During ischemia, the NADH signal initially increased and peaked 5 minutes after ischemia followed by a gradual decline in MIRI groups. NADH fluorescece was significantly lower in HDAC6^-/-^ + MIRI group than in MIRI group 3 to 5 minutes after ischemia and greater in either T1D + MIRI or HDAC6^-/-^ + MIRI groups during a period of 30-min ischemia than in Ctrl group (P < 0.05, n = 8-9 hearts/group) (Figure 6B). During reperfusion, the NADH signal remained relatively stable in the 4 experimental groups. NADH levels were higher in T1D +MIRI groups than MIRI, HDAC6^-/-^ + MIRI, or HDAC6^-/-^ + T1D + HDAC6 groups during the period of 120-min reperfusion (P < 0.05, n = 8-9 hearts/group). No significant differences were found among HDAC6^-/-^ + T1D + MIRI, HDAC6^-/-^ + MIRI and MIRI groups. Thus, HDAC6 KO presevered mitochondrial nicotinamide adenine dinucleotide (NAD^+^) metabolism during ischemia and reperfusion in T1D.

**Figure 6.**
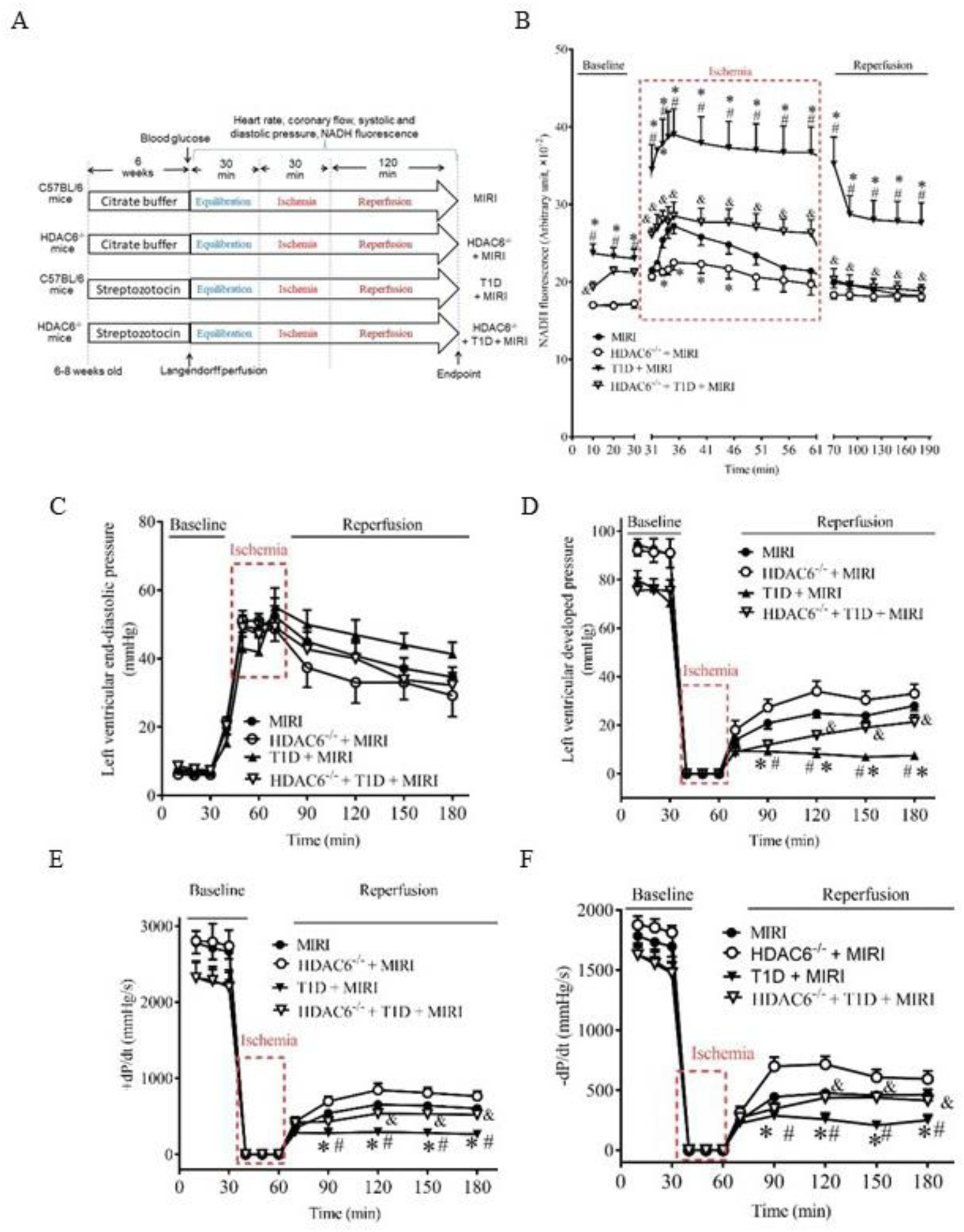
HDAC6 knockout reduced T1D-elicited increases in mitochondrial NADH levels and improved cardiac function during reperfusion in Langendorff-perfused hearts. A: schematic presentation of experimental procedure of *ex vivo* experiments for measurements of NADH and cardiac function of HDAC6^-/-^ and C57BL/6 mice undergoing ischemia/reperfusion injury (MIRI) (n = 8-9 mice/group); B: the dynastic changes in NADH fluorescence of Langendorff-perfused mouse hearts at baseline, ischemia, and reperfusion C: left ventricular end-diastolic function; D: left ventricular developed pressure; E: +dP/dt (maximum rate of increase of left ventricular developed pressure); F: –dP/dt (maximum rate of decrease of left ventricular developed pressure). Data are presented as means ± SEM. Kruskal-Wallis test followed by Dunn’s test was used to analyze multiple group comparisons. *P < 0.05 versus MIRI groups; ^#^P < 0.05 versus HDAC6^-/-^ + MIRI groups, and ^&^P < 0.05 versus T1D + MIRI.

Since NAD^+^/NADH is involved in energy and redox homeostasis,^35^ we determine cardiac function in isolated Langendorff-perfused mouse hearts during ischemia and reperfusion. The dynamic changes in LV pressure and derivatives of HDAC6^-/-^ and C57BL/6 mouse hearts undergoing MIRI are shown in Figure 6. Left ventricular end-diastolic pressure (LVEDP) at baseline was comparable among the 4 groups (P > 0.05) (Figure 6C), whereas the values of left ventricular developed pressure (LVDP) and ±dP/dt at baseline were smaller in T1D + MIRI and HDAC6^-/-^ + T1D + MIRI than MIRI groups (P < 0.05, n = 8 mice/group) (Figures 6D, 6E, and 6F). Global ischemia for 30 minutes resulted in the cessation of the contraction and relaxation of the hearts and an increase in LVEDP. With reperfusion, contraction and relaxation were gradually restored in all mouse hearts. There were no significant differences in LVEDP between T1D + MIRI and MIRI groups during ischemia and reperfusion (P>0.05) (Figure 6C). Compared with MIRI groups, LVEDP was significantly decreased in HDCA6^-/-^ + MIRI groups 1 to 2 h (P < 0.05, n = 8 mice/group) after reperfusion but not in HDAC6^-/-^ + T1D + MIRI groups. The values of LVDP and ±dP/dt were significantly smaller in T1D +MIRI than MIRI groups from 30 minutes to 2 h after reperfusion (P < 0.05) (Figures 6D, 6E, and 6F). Compared with MIRI groups, the values of LVDP and ±dP/dt were significantly increased in HDAC6^-/-^ +MIRI groups from 30 minutes to 2 h after reperfusion and decreased in HDAC6^-/-^+ T1D + MIRI groups (Figures 6D, 6ED and 6F). There were no significant differences between HDAC6^-/-^ + T1D + MIRI and T1D + MIRI groups in the values of ±dP/dt. These results indicate that HDAC6 KO ameliorates cardiac diastolic and systolic dysfunction caused by T2D and MIRI.

### Pharmacological Inhibition of HDAC6 Attenuates Mitochondrial NADH Levels and Improves Cardiac Function in T2D

We examined the effects of HDAC6 inhibition on mitochondrial NADH and cardiac dysfunction during ischemia and reperfusion in T2D and db/+ Ctrl mice. Langendorff-perfused mouse hearts were subjected to 30 min of global ischemia followed by 120 min of reperfusion *ex vivo*. TSA or vehicle was perfused into the heart for 20 min prior to ischemia and continuously throughout 120 min of reperfusion (Figure 7A). NADH fluorescence during 30 min of ischemia was comparable between TSA + MIRI and MIRI groups and much stronger in both T2D + MIRI and TSA + T2D + MIRI groups than MIRI groups (Figure 7B). Compared to the T2D + MIRI group, NADH levels during 30 min of ischemia and 120 min of reperfusion were significantly decreased in TSA + T2D + MIRI group. These results suggest that HDAC6 inhibition potently decreases mitochondrial NADH levels in T2D mice undergoing MIRI.

**Figure 7.**
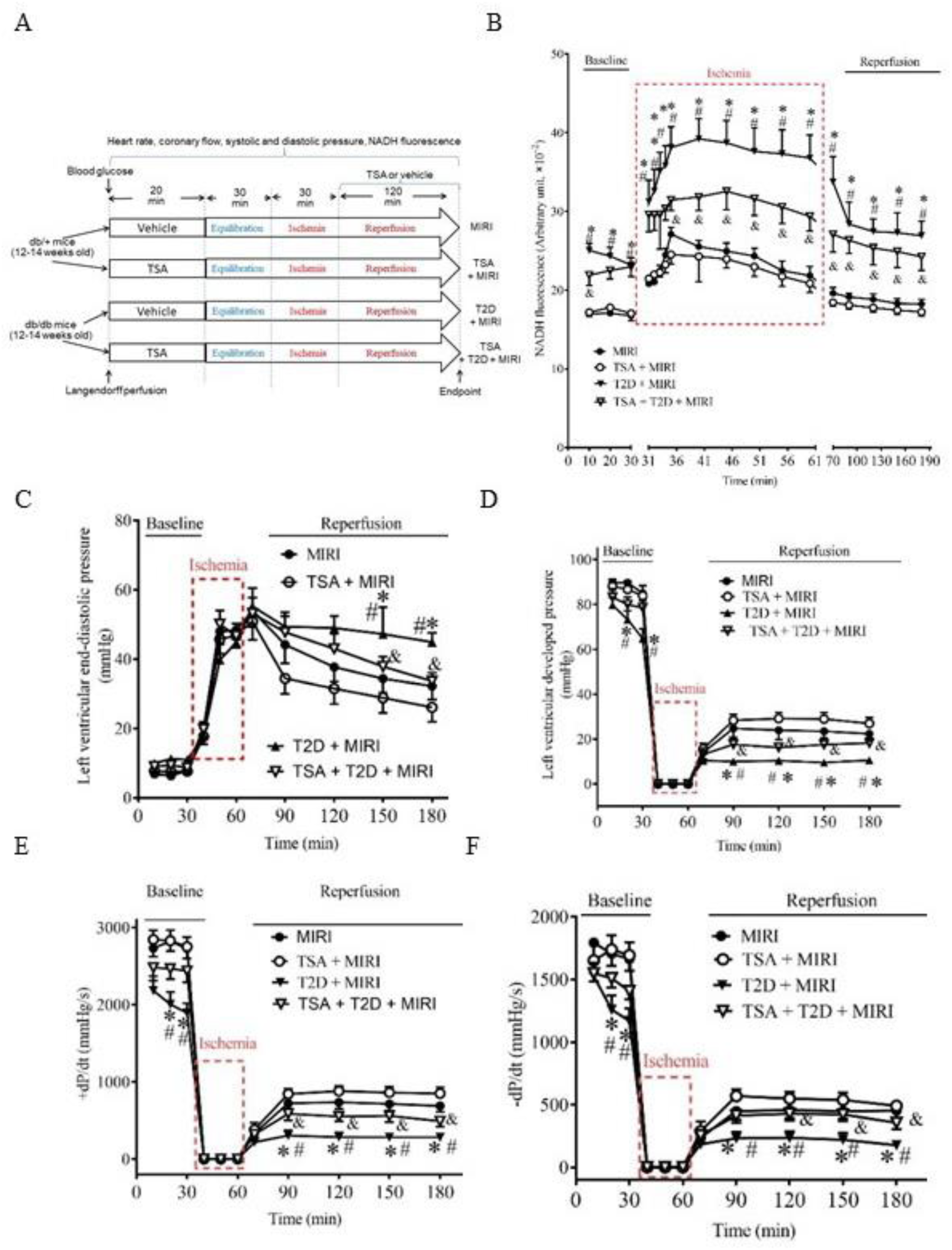
Inhibition of HDAC6 decreased T2D-elicited increases in NADH levels and improved cardiac function during reperfusion in Langendorff-perfused hearts. A: experimental procedures for the measurement of NADH and cardiac function in isolated Langendorff-perfused mouse hearts undergoing global ischemia/reperfusion; B: the dynastic changes in NADH fluorescence of Langendorff-perfused mouse hearts at baseline, ischemia, and reperfusion C: left ventricular end-diastolic function; D: left ventricular developed pressure; E: +dP/dt; F: –dP/dt. Data are presented as means ± SEM. Kruskal-Wallis test followed by Dunn’s test was used to analyze multiple group comparisons. *P < 0.05 versus MIRI groups; ^#^P < 0.05 versus TSA + MIRI groups, and ^&^P < 0.05 versus T2D + MIRI (n = 8-10 hearts/group).

We next quantified LVEDP, LVDP, and ±dP/dt obtained from Langendorff-perfused hearts.^36^ LVEDP at baseline was comparable among the 4 groups (P > 0.05) (Figure 7C), whereas the values of LVDP and ±dP/dt at baseline were smaller in both T2D + MIRI and TSA + T2D + MIRI groups than both MIRI and TSA + MIRI groups (P < 0.05, n = 8 mice/group) (Figures 7D, 7E, and 7F). Global ischemia for 30 minutes resulted in the cessation of the contraction and relaxation of the hearts and an increase in LVEDP. With reperfusion, contraction and relaxation were gradually restored in all mouse hearts. There were no significant differences in LVEDP between TSA + MIRI and MIRI groups during ischemia and reperfusion (P>0.05) (Figure 7C). Compared with MIRI groups, LVEDP was significantly increased in T2D + MIRI groups from 30 minutes to 2 hours (P < 0.05, n = 8 mice/group) after reperfusion but not in TSA + T2D + MIRI groups (n = 9-10 mice/group). The values of LVDP and ±dP/dt were significantly smaller in T2D + MIRI than MIRI groups from 30 minutes to 2 hours after reperfusion (P < 0.05) (Figures 7D, 7E, and 7F). Compared with MIRI groups, the values of LVDP and ±dP/dt were not significantly altered in TSA + MIRI groups and decreased in T2D + MIRI groups from 30 minutes to 2 hours after reperfusion (Figures 7D, 7E, and 7F). All 3 values were greater in TSA + T2D + MIRI than T2D + MIRI groups from 30 minutes to 120 minutes. These results indicate that HDAC6 inhibition improves cardiac diastolic and systolic function in ischemic/reperfused T2D mice.

### HDAC6 Knockdown Blocks the TNFα-induced Inhibition of mCI Activity *in vitro*

Both cardiomyocytes and inflammatory cells generate TNFα which impairs the electron transport of mitochondrial respiratory chain.^21, 37^ To obtain insights into the role of HDAC6 in cardiomyocyte-derived TNFα, we investigated the effects of HDAC6 silencing in H9C2 cardiomyocytes on TNFα levels and mCI activity by cuturing H9C2 cells under 25.0 mM D-dextrose (HG) *in vitro* (Figure 8C). The HDAC6-siRNA significantly decreased HDAC6 mRNA levels and HDAC6 protein expression in H9C2 cells (P < 0.05, n = 5 dishes/group) (Figures 8A and 8B). D-glucose at a concentration of 25.0 mM and hypoxia/reoxygenation together significantly augmented HDAC6 activity, increased TNFα levels, and inhibited mCI activity (Figures 8D-8F). Intriguingly, HDAC6 silencing decreased TNFα and augmented mCI activity of H9C2 cells in HG and HRI.

**Figure 8.**
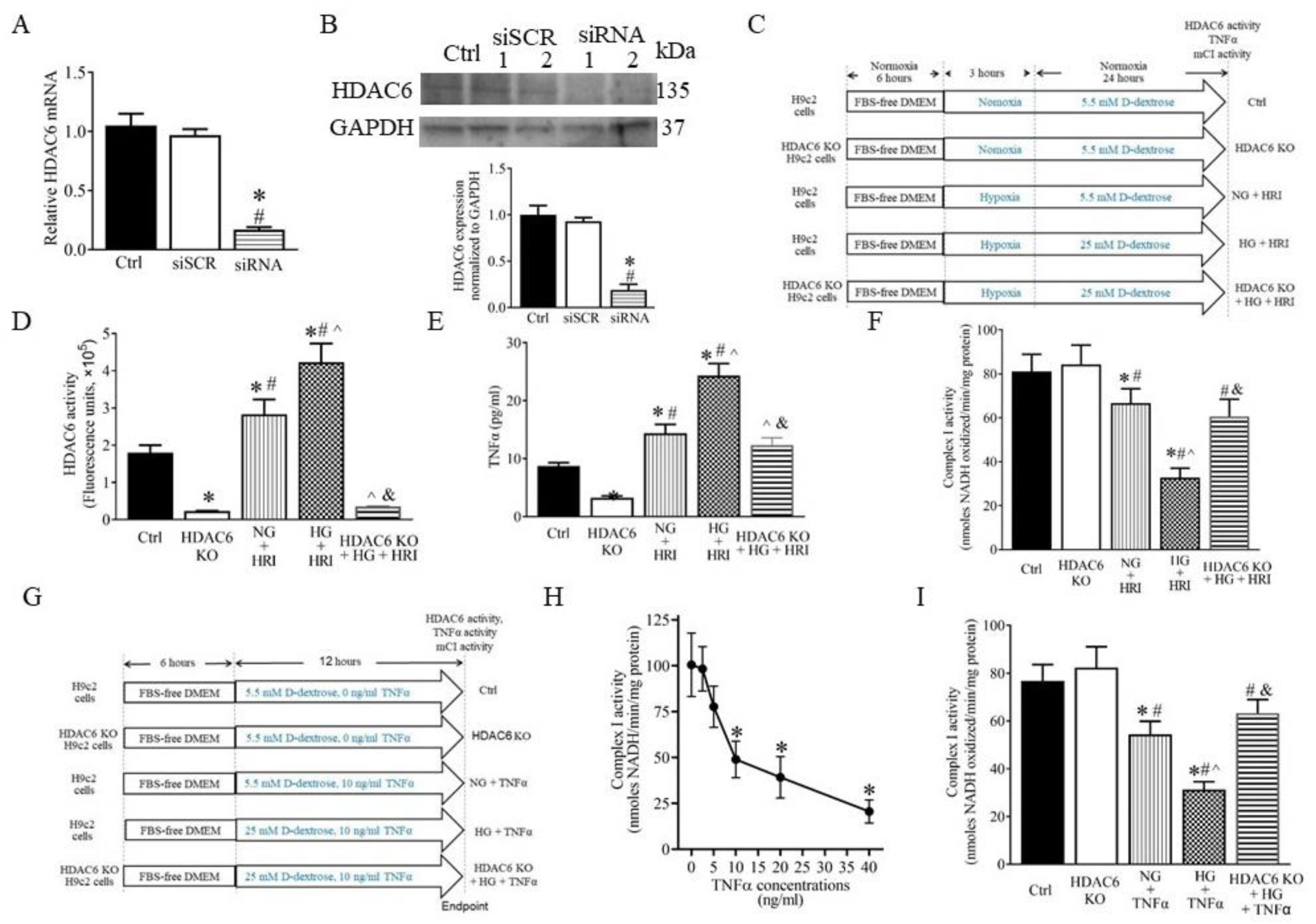
HDAC6 knockdown (KO) blocked TNFα-induced inhibition of mCI activity in cardiomyocytes. A: qRT-PCR analysis of HDAC6 mRNA levels in H9C2 cardiomyocytes; B: Western blot analysis of HDAC6 protein expression in H9C2 cardiomyocytes. Top: representative Western blot bands showing the expression of HDAC6 and GAPDH as control. Bottom: expression of HDAC6 proteins normalized to GAPDH in H9C2 cardiomyocytes. Ctrl = control, siSCR = scrambled siRNA, siRNA 1 = HDAC6-siRNA 1, siRNA 2 = HDAC6-siRNA 2. C: experimental procedures showing the effects of HDAC6 KO on hypoxia/reoxygenation injury (HRI)-induced changes in TNFα and mCI in H9c2 cardiomyocytes; HG = high glucose, NG = normal glucose. D: HDAC6 activity; E: TNFα concentrations; F: mCI activity. *P<0.05 versus Ctrl, ^#^P < 0.05 versus HDAC6 KO, ^^^P < 0.05 versus NG + HRI, ^&^P < 0.05 versus HG + HRI group (n = 9-10 /group). G: experimental procedures showing the effects of HDAC6 KO on HG and exogenous TNFα-induced inhibition of mCI in H9c2 cardiomyocytes; H: dose-dependent effects of TNFα on mCI activity under 25 mg/dl of glucose. *P<0.05 versus 0 ng/ml TNFα groups (n = 6/group). I: HDAC6 KO antagonized TNFα-elicited inhibition of mito CI activity. *P<0.05 versus Ctrl, ^#^P < 0.05 versus HDAC6 KO, ^^^P < 0.05 versus NG + TNFα, ^&^P < 0.05 versus HG + TNFα group (n = 9-10 /group).

Next, we investigated the effects of HDAC6 silencing on exogenous TNFα-induced damage to mCI in H9C2 cardiomyocytes in the presence of 5.5 mM (NG) and 25.0 mM (HG) D-dextrose (Figure 8G). Under 5.5 mM D-dextrose, TNFα from 0-40 ng/ml did not alter mCI activity. However, under 25.0 mM D-dextrose (HG), TNFα dose-dependently decreased CI activity (P < 0.05, n = 6 dishes/group) (Figure 8H). TNFα at 20 ng/ml and 40 ng/ml elicited significant cell necrosis. Since TNFα at 10 ng/ml did not cause cell death, TNFα at a dose of 10 ng/ml was chosen to treat H9C2 cells. Interetingly, 10 ng/ml TNFα-induced decreases in mCI activity were blocked by HDAC6 silencing (P < 0.05 between TNFα + HDAC6 KO and TNFα groups, n = 9-10 dishes/group) (Figure 8I). These data suggest that augmentation of HDAC6 activity increases TNFα production in cardiomyocytes and augmentation of both internal and exogenous TNFα directly inhibits mCI activity.

## DISCUSSION

By dissecting the effects of HDAC6 activity and HDAC6 KO and inhibition on TNFα, mitochondria morphology, mCI, mitochondrial NADH, infarct size, and cardiac function, we found evidence of an essential contribution of HDAC6 in the pathogenesis of MIRI in diabetes. First, by determining HDAC6 activity in T1D and T2D alone and in combination with MIRI, we have found that MIRI and diabetes synergistically augment myocardial HDCA6 activity and TNFα levels and decrease mCI activity. Second, by neutralizing TNFα with an anti-TNFα monoclonal antibody, we have shown that inhibited myocardial mCI activity in ischemic/reperfused diabetic mice is associated with high levels of TNFα. Third, by using HDAC6 KO mice and a selective HDAC6 inhibitor, we demonstrate that HDAC6 inhibition decreased the production of TNFα and NADH and augmented myocardial mCI activity in ischemic/reperfused diabetic mice, along with marked amelioration of cardiac dysfunction. Lastly, in cultured cardiomyocytes, we demonstrated that HDAC6 KO blocked high glucose- and TNFα-elicited suppression of mCI activity. These findings indicate that HDAC6 inhibition protects diabetic hearts against MIRI by reducing TNFα-elicited impairment to mCI in diabetes.

### HDCA6 Regulates Generation of TNFα in Ischemic/Reperfused Diabetic Mice

The deacetylase activity of HDAC6 markedly affects acetylation and deacetylation of histone and nonhistone proteins such as *α*-tubulin, cortactin, and heat shock protein 90.^3, 10, 38^ Augmentation of HDAC6 activity potently increases the expression of TNFα gene by the ROS-MAPK-NF-κB/AP-1 pathway-dependent mechanism in macrophages.^39^ In animal experiments and cultured cardiomyocytes, we have shown that HDAC6 activity and plasma TNFα levels are augmented synergistically by hyperglycemic stress and MIRI/HRI. Moreover, HDAC6 KO or inhibition markedly decreases production of TNFα. Taken together, HDAC6 critically affects TNFα production in ischemic/reperfused diabetic hearts.

Mitochondria do not only coordinate cellular metabolism and regulate calcium homeostasis and ROS production, they are also determinants of cell death.^40^ In response to hyperglycemic and hypoxic stresses, mitochondria undergo fusion and fission. Growing evidence suggests that morphological changes in mitochondria by fusion and/or fission play a critical role in protecting mitochondria from metabolic stresses.^41^ In the current study, we demonstrate that MIRI in diabetic hearts results in excessive production of TNFα and mitochondrial fission, whereas KO or inhibition of HDAC6 reduces plasma TNFα levels and mitochondrial fission. Excessive TNFα induces mitochondrial dysfunction with impaired basal, ATP-linked, and maximal respiration, decreases cellular ATP synthesis, and increases mitochondrial superoxide production.^25^ Collectively, the augmentation of HDAC6 activity in ischemic/reperfused diabetic hearts contributes to mitochondrial dysfunction by augmenting production of TNFα.

HDAC6 KO or inhibition markedly decreased the levels of plasma and myocardial TNFα, not only in non-diabetic mice undergoing MIRI, but also in T1D or T2D subjected to MIRI. Interstingly, in the presence of high glucose, exogeenuous TNFα dose-dependently attenuated the activity of mCI in H9c2 cardiomyocytes. This negative effect of TNFα was decreased by HDAC6 silencing. Previous studies showed that HDAC6 inhibition prevents TNFα-induced caspase 3 activation and α-tublin and β-catenin acetylation.^42, 43^ Therefore, the cardioprotective effects of HDAC6 KO or inhibition is associated with reduction of the TNFα-induced impairments to mCI activity.

### Diabetes and TNFα Impair Mitochondrial Function

The mitochondrial electron transport chain (ETC) is localized in the mitochondrial cristae. The ETC shuttles electrons from NADH and FADH_2_ and transfers them through a series of electron carriers within multiprotein respiratory complexes (complex I to IV) to oxygen, in the process, generating an electrochemical gradient that can be used by the F_1_-F_0_-ATP synthase (complex V) in the mitochondrial inner membrane to synthesize ATP. The respiratory complexes can assemble to form a variety of supercomplexes to prevent ROS production.^44^ However, diabetes impairs mitochondrial supercomplex assembly and activity, which underlies the mechanism of mitochondrial respiratory defects and mitochondrial dysfunction.^45^

In healthy individuals, cardiac TNFα concentrations are low and do not affect mitochondrial function. The presence of TNFα protein and transcripts is usually restricted to microvessels in the normal heart. However, in ischemic/reperfused diabetic hearts, the levels of plasma and myocardial TNFα are highly increased.^46, 47^ Aberrantly high levels of TNFα can impair the mitochondrial supercomplex assembly and activity.^21^ An recent study indicates that TNFα-activated cellular glutamine uptake leads to an increased concentration of succinate, a Krebs cycle intermediate in a zebrafish model of tuberculosis infection.^22^ Oxidation of this elevated succinate by complex II drives RET, thereby generating the mitochondrial ROS superoxide anion at mCI.^22^ Excess TNF in mycobacterium-infected macrophages elevates mitochondrial ROS production by RET through mCI.^22^ In addition, TNFα as a major pro-inflammatory mediator has many unfavorable effects on the heart, including down-regulation of myocardial contractility, ventricular remodeling, increased rate of apoptosis among endothelial cells and myocytes, as well as alteration in the expression and function of the enzymes regulating nitric oxide production.^48^

### Diabetes and MIRI Impair Mitochondrial Supercomplex Assembly and Activity

mCI is composed of 45 subunits including 14 core and 31 supernumerary subunits with lysine residues.^49^ It transfers electrons from NADH to coenzyme Q_10_ (ubiquinone) pumping 4H^+^ into the intermembrane space thereby contributing to the mitochondrial membrane potential.^50^ These electrons are sequentially transferred to Complex III (coenzyme Q-cytochrome c oxidoreductase), cytochrome *c*, and Complex IV (cytochrome *c* oxidase), resulting in the reduction of oxygen to water. By monitoring dynamic changes in mitochondrial NADH levels, we demonstrated abnormal increases in mitochondrial NADH levels of diabetic hearts at baseline and during ischemia, suggesting low activity of mCI in diabetic hearts. Previous studies have shown that diabetes and high levels of TNFα in diabetic hearts impair supercomplex assembly and activity which prevent the production of ROS.^51^ We speculate that in ischemic/reperfused diabetic hearts, high levels of mitochondrial NADH can be attributed to supercomplex defect.

Physiologically, mCI oxidizes NADH to regenerate NAD^+^ to sustain crucial metabolic processes such as the tricarboxylic acid cycle and β-oxidation, and provide electron to the downstream complexes of the electron transport chain.^18^ During MIRI, mCI can also catalyze RET to generate ROS, leading to mitochondrial dysfunction and tissue damage.^52, 53^ Thus, low mCI activity in ischemic/reperfused diabetic hearts decrease not only ATP production which is important for cardiac contraction and relaxation but also the ratio of NAD^+^/NADH.^32, 54^ NAD^+^ and its metabolites function as critical regulators to maintain physiologic processes, enabling the plastic cells to adapt to environmental changes including nutrient perturbation, genotoxic factors, circadian disorder, infection, inflammation and xenobiotics.^55, 56^ Thus, low mCI activity has a profound negative effect on metabolism and function in ischemic/reperfused diabetic hearts.

### HDAC6 is a Therapeutic Target for Cardioprotection Against MIRI in Diabetes

Genetic disruption of HDAC6 did not alter body weight, blood glucose levels, systemic blood pressure, myocardial HDAC6 activity at baseline, and cardiac phenotype and function of mice. The administration of STZ elicited a continuous increase in fasting blood glucose in both C57BL/6 and HDAC6^-/-^ mice. There were no significant differences in blood glucose levels between C57BL/6 and HDAC6^-/-^ mice. Thus, HDAC6 KO does not affect the deleterious effect of STZ on pancreatic β-cells. The present data indicated that either diabetes or MIRI induced an increase in the activity of cardiac HDAC6. These results are consistent with previous studies showing that the activity of cardiac HDAC6 was enhanced in T1D rats or mice.^13, 57^ HDAC6 KO neither markedly affect myocardial area-at-risk and infarct size nor caused changes in LV systolic and diastolic function during MIRI in non-diabetic C57BL/6 mice. Intriguingly, T1D caused increased infarct size and impaired cardiac function in C57BL/6 mice but not HDAC6^-/-^ mice. Emerging evidence suggests that HDAC6 is profoundly implicated in the responses of the heart to MIRI, hypertension, angiotensin II, etc.^4, 14, 15^ Collectively, HDAC6 is an essential negative regulator of MIRI in T1D.

The db/db mouse of leptin receptor deficiency is currently the most widely used model for T2D.^27^ Administration of TSA did not affect myocardial area-at-risk and infarct size in non-diabetic db/+ mice that had undergone MIRI. Interestingly, TSA significantly blocked T2D-induced increase in infarct size. Taken together, the selective inhibition of HDAC6 with TSA protects the heart from MIRI in T2D. To the best of our knowledge, this is the first study to report that HDAC6 KO or inhibiton attenuated MIRI in both T1D and T2D. It is reported that the activity of tissue HDAC6 is markedly elevated in patients with T2D and in db/db mice, concomitant with leptin hyposensitivity.^5^ A recent study showed that the administration of TSA for 2 weeks restores leptin sensitivity and reduces obesity in db/db mice.^3, 5^ Thus, HDAC6 is a critical component of T2D hearts in response to MIRI.

TEM examination of the ultrastructure of the myocardium in mouse hearts can provide visual evidence of mitochondria. Both T1D and T2D resulted in the exacerbation of mitochondrial swelling and dissolution in ischemic/reperfused myocardium. It is reported that hyperglycemia in diabetes mellitus increases the production of ROS during MIRI.^58^ Excessive ROS can impair mitochondrial structure and function.^59^ We speculate that excessive production of ROS in diabetes contributes to more severe impairment of mitochondrial structure. Interestingly, genetic disruption or inhibition of HDAC6 preserved mitochondrial structure during MIRI in diabetes mellitus. Accumulating evidence suggests that HDAC6 is involved in redox regulation and cellular stress responses.^60^ Myocardial HDAC6 activity is jointly augmented by diabetic stressed and MIRI.^13, 57^ Taken together, HDAC6 is a promising therapeutic target for reducing oxidative stress in diabetic myocardium.

Importantly, either genetic disruption or inhibition of HDAC6 markedly attenuates MIRI in the mice with T1D and T2D. The cardioprotective effects of HDAC6 knockout or inhibition are associated with decreased TNFα levels and preservation of mCI activity. Interestingly, HDAC6 KO directly blocks TNFα-elicited impairments to mCI in cardiomyocytes. These findings support the notion that HDAC6 can serve as a therapeutic target for the protection of diabetic hearts against MIRI.

In summary, the current study demonstrates that either T1D or T2D and MIRI synergistically augment HDAC6 activity and production of TNFα. Aberrant increases in TNFα suppress myocardial mCI activity, leading to increases in NADH levels during myocardial ischemia in diabetes. These negative changes impair cardiac systolic and diastolic function. HDAC6 knockout or inhibition not only decreases the production of cardiac TNFα but also blocks TNFα-induced impairments to mCI. This study suggests that HDAC6 is an essential negative regulator of MIRI in diabetes.

RI and cardiac function in diabetes. Selective inhibition of HDAC6 has high therapeutic potential for acute IHS in diabetes.

## Nonstandard Abbreviations and Acronyms

Cox I: mitochondrial cytochrome c oxidase I
Ctrl: control
DMEM: Dulbecco’s modified Eagle’s medium
DMSO: dimethylsulfoxide
+dP/dt: maximum rate of increase of left ventricular developed pressure
-dP/dt: maximum rate of decrease of left ventricular developed pressure
EDTA: ethylenediaminetetraacetic acid
ELISA: enzyme-linked immunosorbent assay
FBS: fetal bovine serum
GAPDH: glyceraldehyde-3-phosphate dehydrogenase
HG: high glucose
HRI: hypoxia/reoxygenation injury
IHS: ischemic heart disease
KDEL: Lys-Asp-Glu-Leu
KO: knockout
HDAC: histone deacetylase
LV: left ventricle
LVDP: left ventricular developed pressure
LVEDP: left ventricular end-diastolic pressure
MIRI: myocardial ischemia/reperfusion injury
mCI: mitochondrial complex I

